# Engineered *Acinetobacter baylyi* ADP1-ISx cells are sensitive DNA biosensors for antibiotic resistance genes and a fungal pathogen of bats

**DOI:** 10.1101/2025.05.17.654686

**Authors:** Jeffrey Chuong, Keaton W. Brown, Isaac Gifford, Dennis M. Mishler, Jeffrey E. Barrick

**Affiliations:** Department of Biomedical Engineering, The University of Texas at Austin, Austin, TX 78712, USA; Department of Molecular Biosciences, Center for Systems and Synthetic Biology, The University of Texas at Austin, Austin, TX 78712, USA; The Freshman Research Initiative, College of Natural Sciences, The University of Texas at Austin, Austin, Texas, USA

**Keywords:** Natural transformation, Cell-based biosensor, Environmental DNA, Antibiotic resistance genes, White-nose syndrome, Biosurveillance

## Abstract

Naturally competent bacteria can be engineered into platforms for detecting environmental DNA. This capability could be used to monitor the spread of pathogens, invasive species, and resistance genes, among other applications. Here, we create *Acinetobacter baylyi* ADP1-ISx biosensors that detect specific target DNA sequences through natural transformation. We tested strains with DNA sensors that consisted of either a mutated antibiotic resistance gene (TEM-1 *bla* or *nptII*) or a counterselectable gene flanked by sequences from the fungus *Pseudogymnoascus destructans*, which causes white-nose syndrome in bats. Upon uptake of homologous DNA, recombination restored antibiotic resistance gene function or removed the counterselectable gene, enabling selection of cells that sensed the target DNA. The antibiotic resistance gene and *P. destructans* biosensors could detect as few as 3,000 or 5,000,000 molecules of their DNA targets, respectively, and their sensitivity was not affected by excess off-target DNA. These results demonstrate how *A. baylyi* can be reprogrammed into a modular platform for monitoring environmental DNA.

---

The bacterium *Acinetobacter baylyi* has powerful and flexible genetic capabilities for synthetic biology^1,2^. Foremost among these is its ability to efficiently integrate foreign DNA into its chromosome through natural transformation^3–5^, which has been used for genome streamlining^6,7^, metabolic engineering^8–10^, and directed evolution^11^ studies. *A. baylyi* also has potential as a cell-based biosensor for applications in bioremediation and environmental monitoring. It has been engineered to detect pollutants such as salicylate, toluene, and xylene using transcriptional reporters^12,13^. *A. baylyi’s* high natural competence has also been used to create sensors for extracellular DNA, including detecting DNA from colorectal cancer cells *in vivo*^14^, ancient DNA from a woolly mammoth bone^15^, and antibiotic resistance genes in transgenic plants and soil^16–18^. Here, we design and test two types of *A. baylyi* cell-based DNA biosensors for environmental DNA sequences.

Antibiotic resistance genes (ARGs) have emerged as a threat to global public health due to widespread use of antibiotics in medicine and agriculture^19^. Horizontal gene transfer spreads ARGs among bacterial communities in soil^20,21^ and multidrug-resistant pathogens in hospitals^22^. Monitoring ARGs is necessary to guide interventions to reduce their prevalence. To measure how effectively *A. baylyi* could detect ARGs in environmental DNA, we constructed two biosensor strains, one for a *bla* gene and one for an *nptII* gene. Each was created by inserting a mutated, nonfunctional version of the ARG into the chromosome of the transposon-free *A. baylyi* ADP1-ISx strain^6^. The TEM-1 *bla* gene encodes a β-lactamase enzyme that inactivates β-lactam antibiotics^23,24^, including ampicillin. The *nptII* gene encodes a phosphotransferase enzyme that inactivates aminoglycoside antibiotics, including kanamycin^25^. Each inserted gene was inactivated by introducing a 10-base-pair frameshifting deletion^16^ (**Table S1**). When *A. baylyi* biosensor strains are cultured in the presence of DNA containing their target ARG, some cells will import it and integrate it into their chromosomes through homologous recombination, repairing the inactivated ARG. Cells that have successfully detected the target DNA in this way can be enumerated on selective plates containing the respective antibiotic (**Figure 1a**).

**Figure 1.**
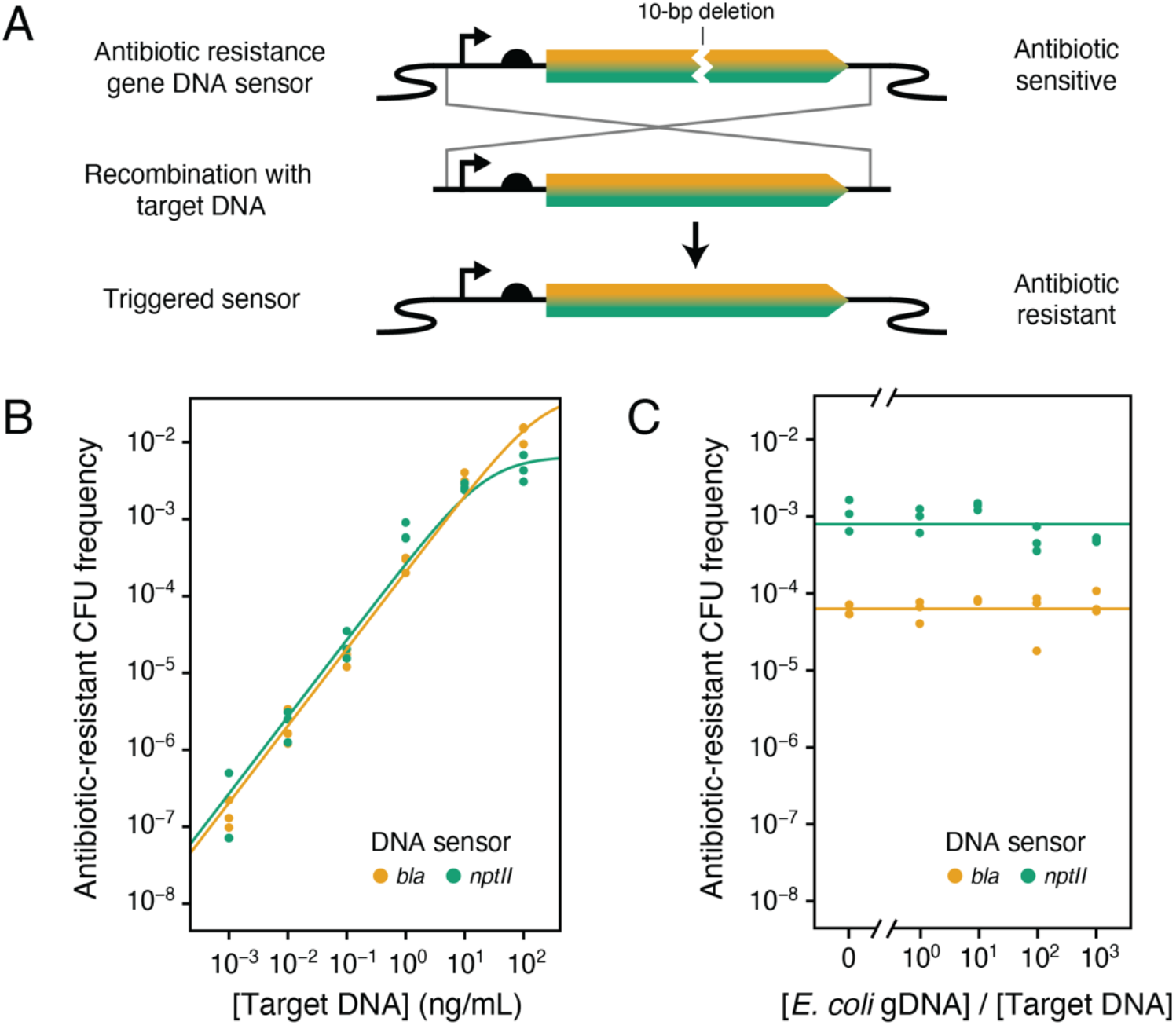
Detection of antibiotic resistance genes. **(A)** Design of *A. baylyi* biosensor strains for detecting the DNA sequences of the *TEM-1 bla* and *nptII* antibiotic-resistance genes. After cells import DNA containing the target ARG through natural competence, homologous recombination will repair the 10-base pair deletion in the ARG sensor in the chromosome, resulting in a functional resistance gene. Plating on the respective antibiotic allows for the selection of cells that have detected the target DNA. **(B)** Frequencies of antibiotic-resistant colony-forming units (CFUs) after transformation with a PCR product of the intact *TEM-1 bla* or *nptII* DNA sequence sensed by each strain. Data points are for biological replicates. The curve is a fit to a hyperbolic model that saturates after maintaining a log-linear relationship at low target DNA concentrations. **(C)** Frequencies of antibiotic-resistant CFUs after transformation with a mixture consisting of 1 ng/mL of target DNA PCR product and off-target *E. coli* genomic DNA at the specified relative concentrations. Data points are for biological replicates. Horizontal lines are the mean of all measurements for each sensor.

We quantified the sensitivity of each *A. baylyi* antibiotic resistance biosensor by adding different amounts of on-target ARG PCR product and measuring how this changed the frequency of antibiotic-resistant cells after allowing time for natural transformation during growth. As the target DNA concentration increased 10-fold, the number of resistant cells increased 10-fold, and this linear relationship was maintained until saturating at around a 10^−2^ frequency of resistant cells in the population when >10 ng/mL target DNA was added (**Figure 1b**). According to the linear relationship between target DNA concentration and resistance frequency, observing one resistant colony in 500 µL of culture containing 10^9^ *A. baylyi* cells would correspond to an estimated detection limit of 6000 molecules of the TEM-1 *bla* gene or 3100 molecules of the *nptII* gene, calculated from the extrapolated DNA concentration that would have a 95% chance of yielding at least one transformant according to a binomial test. If spontaneous mutations could reactivate the ARG sensor, it would result in the detection of false positives, but this rate is extremely low: no resistant colonies were observed in controls without target DNA. We next sought to determine whether the ARG biosensors could detect the target DNA sequence in a mixture containing an excess of off-target DNA. Adding increasing amounts of *E. coli* genomic DNA, ranging from an equal number of total base pairs to the ARG DNA target PCR product up to 1000-fold times as much, did not affect the transformation frequency of the ARG biosensor strain (*p* = 0.39 for the *bla* biosensor, *p* = 0.11 for the *nptII* biosensor, two-tailed Welch’s *t*-test comparing log-transformed frequencies with no added *E. coli* DNA to those 1000-fold as much) (**Figure 1c**). Thus, *A. baylyi* biosensors can efficiently search through large amounts of DNA to recognize a specific gene sequence.

Based on these results, we next sought to detect the presence of a pathogen using a more flexible sensor that recognizes a target DNA sequence that does not encode a selectable function. *Pseudogymnoascus destructans*, the fungus that causes white-nose syndrome in bats, has led to catastrophic declines in North American bat populations, disrupting ecosystems that rely on bats for insect control and pollination^26–28^. The fungus can persist in soils, guano, and cave substrates for extended periods^29^. Early detection of sites contaminated with *P. destructans* could help mitigate its spread. We created a sensor for the DNA sequence of a specific Ty3 long terminal repeat (LTR) retrotransposon family (**Table S2**). There are 82 copies of this retrotransposon in the *P. destructans* genome (**File S1**), and no matches to its sequences were found in the genome of the related species *Pseudogymnoascus pannorum*. The retrotransposon sensor contains a dual-selection *tdk/kanR* cassette^3,6^, which confers azidothymidine (AZT) susceptibility and kanamycin resistance, flanked with homology arms matching the *P. destructans* retrotransposon. When the target sequence is imported into a biosensor cell and integrates into its chromosome, the *tdk/kanR* cassette is deleted, allowing the cell to grow on selective plates containing AZT (**Figure 2a**).

**Figure 2.**
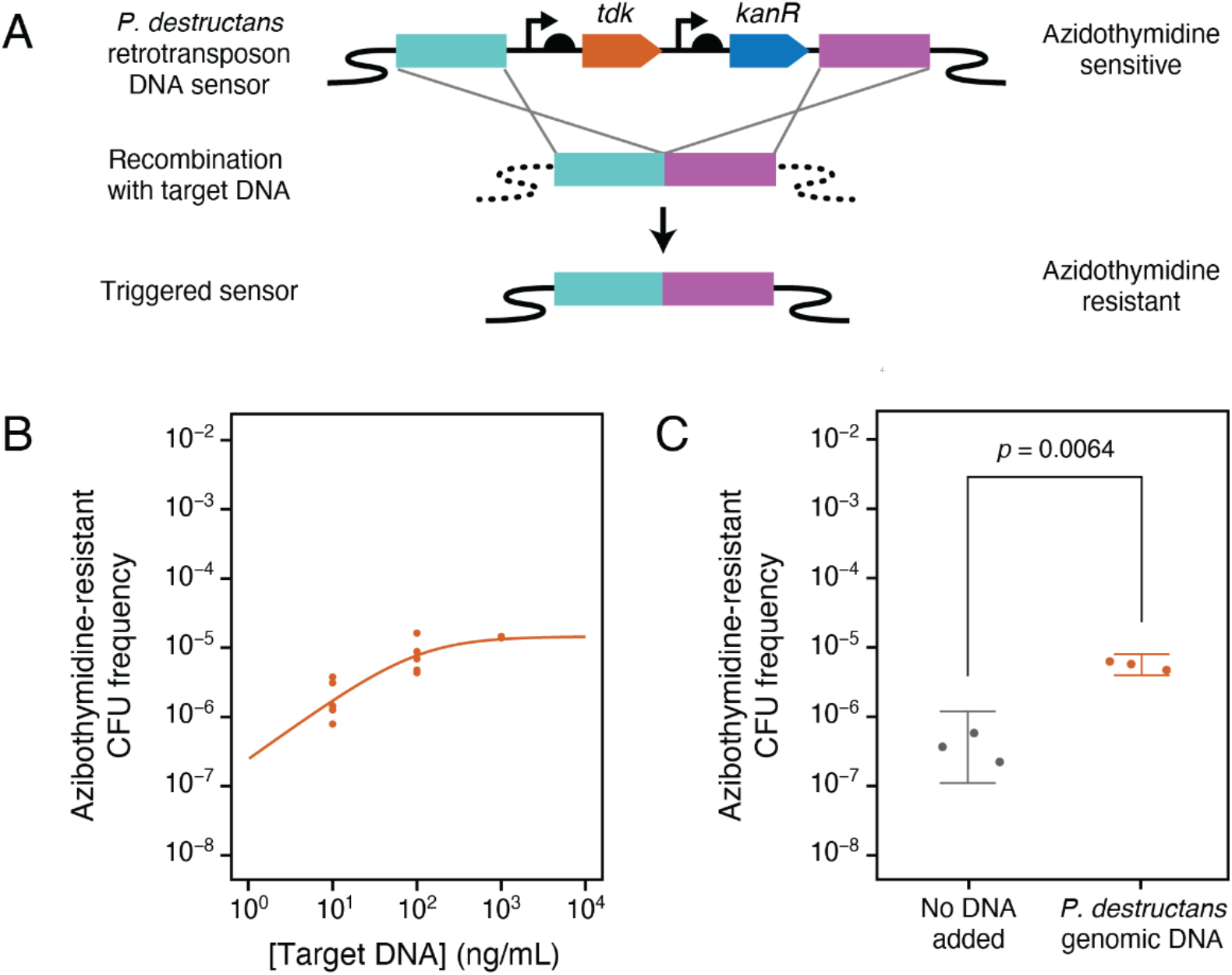
Detection of *P. destructans* DNA. **(A)** Design of the *A. baylyi* biosensor for detecting the DNA sequence of a Ty3 LTR retrotransposon family found in the *P. destructans* genome. Uptake through natural competence and integration of the target DNA sequence into the *A. baylyi* chromosome through homologous recombination removes the *tdk/kanR* cassette, enabling growth in the presence of azidothymidine (AZT). **(B)** Frequencies of AZT-resistant CFUs after transforming the biosensor strain with increasing concentrations of purified PCR product of the *P. destructans* retrotransposon target sequence. The curve is a fit to a hyperbolic model. **(C)** Frequencies of AZT-resistant CFUs when no DNA is added and after transformation with 1.1 µg/mL *P. destructans* total genomic DNA. Background azidothymidine resistance when no DNA is transformed occurs when mutations inactivate the *tdk* gene. The log-transformed frequency of AZT-resistant cells was significantly higher than the background mutation rate when *P. destructans* total genomic DNA was present (two-tailed Welch’s *t*-test).

To quantify the DNA detection capabilities of the *P. destructans* biosensor, we added increasing amounts of retrotransposon PCR product to cultures, allowed time for transformation during growth, and measured the final frequencies of AZT-resistant cells. The frequency of resistant cells scaled close to linearly with target DNA amount at lower concentrations (**Figure 2b**). However, successful detection required DNA concentrations of at least 10 ng/mL, and the frequency of resistant cells in the presence of high DNA concentrations saturated at around 10^−4^. This reduced performance compared to the ARG biosensors is likely due to how incoming DNA must replace the larger *tdk/kanR* cassette for detection by this biosensor versus templating the insertion of just 10 base pairs. Since the *tdk* gene can spontaneously mutate, leading to false positives, we further evaluated the detection limit of the *P. destructans* biosensor by measuring the background *tdk* inactivation rate. We also aimed to examine whether the biosensor could successfully detect the retrotransposon in *P. destructans* genomic DNA, given that all copies of this target sequence together are only 0.5% of its overall sequence. The AZT-resistant colony frequency when the biosensor strain was transformed with 1.1 µg of *P. destructans* genomic DNA was higher than the frequency of *tdk* inactivation by mutations (*p* = 0.005, two-tailed Welch’s *t*-test on log-transformed frequencies) (**Figure 2c**). Given this background and the fit to the results of detecting the PCR product, we estimate that this sensor could detect as few as 5.2 million copies of the *P. destructans* genome in a 500 µl culture with 95% confidence (one-tailed *t*-test on log-transformed frequencies).

In summary, we demonstrated two ways that *A. baylyi* can be engineered to detect specific DNA sequences in environmental DNA using its innate competence machinery and homologous recombination capabilities. The sensor used to detect *P. destructans* DNA was constructed from modular parts using a standardized system of Golden Gate Assembly (GGA) overhangs^30,31^. This design could be easily adapted to detect any DNA sequence of interest, provided the target is long enough to allow for efficient homologous recombination. In the future, several modifications could increase the sensitivity and specificity of DNA-sensing *A. baylyi*. Expression of phage recombinases, such as a RecET system that has been shown to function in *Acinetobacter baumanii*^32^, could increase recombination rates and reduce homology-length requirements. More complex sensing circuits that incorporate transcriptional regulators, fluorescent protein reporters, and/or toxin-antitoxin systems for additional selection and counterselection guards could reduce false positive predictions resulting from spontaneous mutations^33,34^. Lastly, *A. baylyi*’s native CRISPR-Cas system is active^7,35^ and could be programmed to provide extra discrimination against closely related, off-target sequences, such as related fungal pathogens, by targeting their sequences for cleavage^34^.

Recently, the naturally competent bacterium *Bacillus subtilis* has also been engineered as a DNA biosensor for multiplex identification of pathogenic bacteria^33^ and to detect human DNA sequences with different single-nucleotide polymorphisms^34^. Enzyme-based detection assays that rely on DNA amplification generally require input samples to be free of inhibitors that are common in soil and tissue samples^36–38^. By contrast, cell-based *A. baylyi* and *B. subtilis* biosensors should be able to detect DNA in any environment in which they remain viable and competent, which could include *in situ* detection in soil or water samples. However, target DNA molecules must be released from an organism and may be quickly fragmented and degraded in the environment^39,40^. Thus, *A. baylyi* biosensors like those developed here should be further tested and optimized under field conditions, which could include soil samples from bat conservation sites^27,41^ or water samples from wastewater treatment plants^42^, to implement cell-based environmental DNA surveillance.

## Methods

### Culture and Transformation Conditions

*A. baylyi* strains were cultured in the Miller formulation of Lysogeny Broth (LB) at 30°C. Overnight cultures were grown in 3 mL of LB in 18 × 150 mm glass test tubes. For transformations, 35 µL of overnight *A. baylyi* culture and 20 µl of the target DNA was added to 500 µL LB media and incubated for 16 hours at 30°C with shaking at 200 r.p.m. Golden Gate Assembly DNA, purified PCR products, or genomic DNA was used as the target DNA. Transformations were plated on LB agar plates supplemented with 50 µg/mL Kanamycin (Kan), 100 µg/mL ampicillin (Amp), or 200 µg/mL azidothymidine (AZT) for selection or counterselection.

### Biosensor Strain Construction

Engineered strains were created from transposon-free *Acinetobacter baylyi* ADP1-ISx^6^ using Golden Transformation^7^. Sequences of primers and DNA sensor constructs are provided in **Table S1**. For the β-lactamase ARG biosensor, a non-functional version of the TEM-1 *bla* gene was inserted into the ADP1-ISx chromosome in place of the *acrB* gene. For the kanamycin ARG biosensor, a non-functional version of the *nptII* gene was inserted into the chromosome in place of the *ACIAD2049* gene. Template DNA for amplifying the *acrB* and *ACIAD2049* homology flanks was isolated from *A. baylyi* ADP1-ISx using a PureLink Genomic DNA Mini Kit (Invitrogen). Primers were designed to PCR amplify ARG fragments and then full constructs that contained deletions of 10-base pairs. These pieces were amplified from pKD13 (*nptII*, GenBank AY048744) or pBTK404 (TEM-1 *bla*) plasmid DNA purified using a QIAprep Spin Miniprep Kit (Qiagen). PCR was performed using Phusion™ High-Fidelity DNA Polymerase (New England Biolabs), and products were purified using a DNA Clean & Concentrator-5 kit (Zymo Research). For the *P. destructans* biosensor strain, roughly 2000 base pairs were selected from the consensus sequence of the target Ty3 LTR retrotransposon family and divided into two 1000-base-pair homology flanks. This retrotransposon family was identified by running RepeatModeler^43^ and RepeatMasker^44^ on the *P. destructans* 20631-21 genome (GenBank: GCF_001641265.1). Its sequence conservation was evaluated by aligning the nucleotide sequences of the 82 matches with Muscle. BLASTN searches against the *Pseudogymnoascus pannorum* ATCC 16222 genome (GenBank: GCA_001630605.1) did not identify any similar elements. Sensor homology flanks were PCR amplified from *P. destructans* 20631-21 genomic DNA (American Type Culture Collection MYA-4855D) and extracted from a 1% agarose gel using the Zymoclean Gel DNA Recovery Kit (Zymo Research).

Primers were designed with BsmBI and/or BsaI recognition sites for cloning into an entry vector plasmid and/or ligation into a linear DNA product for use in a standard two-step *A. baylyi* Golden Transformation genetic engineering workflow^7^ based on assembly of YTK/BTK parts^30,31^. Briefly, the Type 2-4 *tdk/kanR* cassette part from pBTK622 was ligated to a roughly 1000-base pair 5′- and 3′-homology regions flanking either the *ACIAD2049* or *acrB* genes in the *A. baylyi* chromosome PCR-amplified as Type 1 and Type 5 parts, respectively. For the antibiotic resistance biosensor strains, PCR products of each mutated ARG, were amplified as Type 2-4 parts and ligated to the same homology flanks for integration into the *A. baylyi* chromosome. For the *P. destructans* biosensor, purified PCR products of the 5′ and 3′ sensor homology flanks matching the *P. destructans* retrotransposon (amplified as Type 1 and 5 parts, respectively) were ligated to PCR products of the 5′ and 3′ homology flanks of *ACIAD2049* (amplified as Type 8 and Type 6 parts, respectively). Homology flanks were ligated together and cloned into an entry vector as composite parts before a final three-piece Golden Transformation that added these flanks to each side of the *tdk/kanR* cassette from pBTK622. *A. baylyi* was cultured and transformed as indicated above to construct the sensor strains.

The genome sequences of each *A. baylyi* DNA biosensor strain were validated using Nanopore sequencing. High molecular weight DNA was extracted with a Quick-DNA HMW MagBead kit (Zymo Research), and sequencing was carried out on a MinION Mk1C sequencer (Oxford Nanopore Technologies). Genome assemblies were generated using Flye^45^ and aligned to the designed sensor constructs to confirm that they were complete and unmutated.

### DNA Detection Assays

For detection of purified PCR products, DNA concentrations ranging from 1 pg/mL to 100 ng/mL were added to transformation cultures, as described above. For detection of *P. destructans* genomic DNA, 1.1 µg/mL of genomic DNA was used. Controls with no added DNA were also tested. After transformation, cultures were serially diluted and spot plated on LB-Kan, LB-Amp, or LB-AZT. Transformations were also plated on LB agar plates with no selective agents to determine total CFU counts. Each DNA detection assay was performed with 3-5 biological replicates. Raw counts and calculated frequencies of resistant colonies are provided in **Table S2** for all assays.

Following prior work^34,46^, frequency measurements of resistant CFUs were fit to a hyperbolic curve with the equation: log { *f* (c) } = log { *f*_max_ [DNA] / (*c*_0.5_ + [DNA]) }. In this equation, *c* is the target DNA concentration, *f* (*c*) is the frequency of resistant CFUs at a given DNA concentration, *f*_max_ is the maximum frequency at saturating DNA concentrations, and *c*_0.5_ is the DNA concentration that yields half the maximum CFU frequency. Detection limits were estimated as the lowest DNA concentration in which one resistant colony would be expected to appear when plating all cells in a 500 µL culture of *A. baylyi*. Model fitting and two-tailed Welch’s *t*-tests comparing log-transformed resistant colony frequencies were performed in R (v4.2.2).

## Supporting Information

**Table S1**. Sequences of primers and sensor constructs.

**Table S2**. Data for all DNA sensing assays.

**File S1**. Alignment of all copies of the target Ty3 LTR retrotransposon family found in the *P. destructans* genome in FASTA format.

## Acknowledgements

We thank Vivek Beeram, Nate Brant, Adam Franco, Macy Horn, Leia Jiang, Sandy Nguyen, Samer Salman, Sai Senapathi, Bill Tang, and Neil Tian for assistance with project planning and performing initial experiments during their participation in the 2022 UT Austin iGEM team; and Daniel Deatherage for assistance with genome sequencing. This work was funded by the National Science Foundation (MCB-2123996 and IOS-2103208 to J.E.B.). The University of Texas at Austin College of Natural Sciences and Department of Molecular Biosciences provided additional support for iGEM participation.

